## Supplemental Figures and Text for "N6-methyladenosine in DNA promotes genome stability"

**Supplemental Figure Legends**

*Figure S1. Generation of mCherry-tagged UNG KO DLD-1 cells.*

(A) Percentage of reads from sequencing of DLD-1 cells after gene editing corresponding to gene transcripts that are unmodified, modified in in-frame, contain a frameshift, or are noncoding. (B) Representative immunoassay using the Simple Western Jess system showing protein expression levels of UNG in two HPRT-targeted and two UNG-targeted clones from A. $\alpha$-tubulin represents loading control. (C) Representative immunoblot showing protein expression levels of UNG2-mCherry constructs expressed in UNG KO DLD-1 cells. $\alpha$-tubulin represents loading control.

*Figure S2. UNG2 responds to uracil-based DNA damage in a manner partly dependent on its intrinsically disordered region.*

(A) Percentage of EdU+ cells in indicated DLD-1 cells. Cells were pulsed with 10 $\mu$M EdU prior to fixation and staining with Invitrogen Click-It$^{TM}$ Alexa Fluor 488 Kit. HPRT indicates WT cells. These cells were targeted with a cutting control targeting to the intronic region of HPRT gene. (B) Representative images from DLD-1 UNG KO cells expressing indicated mCherry-tagged cDNAs upon treatment with increasing concentrations of RTX at 64 hours post-treatment. (C) Quantification of experiment represented in A for percentage of cells with >5 mCherry foci. Error bars, mean ± SEM; Ordinary one-way ANOVA with Dunnett's multiple comparisons test with a single pooled variance, $**p \leq 0.01$, n = 3 biological replicates. Statistical tests performed within individual groups, EV or UNG2, respectively. (D) Representative images of mCherry staining in DLD-1 UNG KO cells expressing UNG2 or EV constructs upon treatment with indicated compounds at 64 hours. (E) Quantification of experiment represented in D for percentage of cells with >5 mCherry foci. Error bars, mean ± SEM; Ordinary one-way ANOVA with Dunnet's multiple comparisons test with a single pooled variance, $**p \leq 0.01$, n = 3 biological replicates.

(F) Representative images of yH2AX staining in DLD-1 UNG KO cells expressing UNG2-mCherry constructs upon treatment with indicated compounds at 64 hours from same experiment as D, E. (G) Quantification of experiment represented in F for average mean nuclear intensity. Error bars, mean ± SEM; Ordinary one-way ANOVA with Tukey's multiple comparisons test with a single pooled variance, ∗∗∗p ≤ 0.01, ∗∗∗p ≤ 0.001, n = 3 biological replicates for all except HU where n=2 biological replicates. (H) MTS cell proliferation in DLD-1 UNG KO cells expressing indicated mCherry-Cry2-tagged cDNAs upon treatment with floxuridine at increasing concentrations. Error bars, mean ± SEM. n=4 biological replicates. (I) KEGG pathway analysis for genes that sensitize cells to floxuridine with log2(foldchange) >|0.5| and -$\log_{10}$(p-value) >2.

RTX, raltitrexed; Flox, floxuridine; EV, empty vector; IDR, intrinsically disordered region; FDR, false discovery rate

*Figure S3. N6-methyladenosine foci in response to genomic uracil-inducing agents are not RNA:DNA hybrids.*

(A) MTS cell proliferation in SW620 cells upon treatment with 15 μM METTL3 inhibitor and indicated compounds. Error bars, mean ± SEM, n=3, technical replicates, representative of 3 biological replicates. Representative of 3 biological replicates. (B) Representative immunoblot images from DLD-1 cells nucleofected with ribonucleoproteins containing Cas9 and indicated guide RNAs (gRNA) as performed for C. Antibodies used for blotting indicated on the left. α-tubulin represents loading control. (C) Growth curves in DLD-1 cells upon treatment of floxuridine. Error bars, mean ± SD, n=3 technical replicates. (D) Representative images of immunofluorescence staining with anti-N6-methyladenosine antibody in DLD-1 cells after fixation and permeabilization, but without pre-extraction in the presence or absence of floxuridine. (E) Agarose gel showing 1 μg of RNA extracted from whole cells with or without treatment with RNase A for 1 hour prior to loading on gel. (F) Representative images from DLD-1 cells upon treatment with floxuridine or raltitrexed for 66 hours. Prior to staining with anti-N6-methyladenosine antibody, indicated samples were treated with RNase H. (G) Quantification of experiment represented in F for percentage of cells with >10 N6-methyladenosine foci. Error bars, mean ± SEM; Ordinary one-way ANOVA with Tukey's multiple comparisons test with a single pooled variance, $*p \leq 0.05$, $****p \leq 0.0001$, n = 3 biological replicates for floxuridine treatment and n=4 biological replicates for DMSO and raltitrexed treatment. (H) Quantification of mean nuclear DAPI staining from experiment in Figure 3D, E. Error bars, mean ± SEM; Krustal Wallace with Dunn's multiple comparisons test, $**p \leq 0.01$, n = 5 biological replicates. (I) Representative images from DLD-1 cells upon treatment with raltitrexed for 66 hours. Prior to staining with anti-N6-methyladenosine antibody, indicated samples were treated with RNase A or DNase. (J) Quantification of experiment represented in I for percentage of cells with >5 N6-methyladenosine foci. Error bars, mean ± SEM; RM one-way ANOVA with Dunnet's multiple comparisons test with a single pooled variance, $***p \leq 0.001$, n = 4 biological replicates. (K) Representative images from DLD-1 cells upon treatment with raltitrexed for 66 hours in the presence or absence of METTL3 inhibitor. (L) Quantification of experiment represented in K for percentage of cells with >10 6mA foci. Error bars, mean ± SEM; Ordinary one-way ANOVA with Dunnet's multiple comparisons test with a single pooled variance, $*p \leq 0.05$, $***p \leq 0.001$, n = 4 biological replicates.

Flox, floxuridine, RTX, raltitrexed

*Figure S4. The presence of 6mA does not alter UNG binding kinetics.*

(A) Representative immunoblot of whole cell lysates from DLD-1 cells in the presence or absence of METTL3 inhibitor blotted with UNG or β-actin antibodies. β-actin represents loading control. (B) Equilibrium dissociation constants ($K_d$) measured by biolayer interferometry for binding of UNG to indicated dsDNA templates. Mean ± SEM for n=3 biological replicates displayed in table. (C) Equilibrium dissociation constants ($K_d$) measured by biolayer interferometry for binding of UNG to indicated ssDNA templates. Mean ± SEM for n=2 biological replicates displayed in table.

*Figure S5. 6mA foci, but not UNG2 foci, correlate with DNA damage levels.*

(A) Representative images of yH2AX staining in DLD-1 cells upon treatment with indicated compounds at 64 hours from same experiment as Figure 5 D, E. (B) Quantification of experiment represented in A for average mean nuclear intensity. Error bars, mean ± SEM; Ordinary one-way ANOVA with Tukey's multiple comparisons test with a single pooled variance, ***$p \leq 0.01$, ***$p \leq 0.001$, n = 3 biological replicates for all except HU where n=2 biological replicates. (C). Co-staining with anti-mCherry and anti-N6-methyladenosine antibodies in cells treated with indicated drugs. Green arrows indicate cells where both mCherry and 6mA staining are present and white arrows indicate cells where only 6mA staining is present.

Flox = floxuridine, RTX = raltitrexed, HU = hydroxyurea, Gem = gemcitabine, MMC = mitomycin

*UPLC Mass Spectrometry*

Mass spectrometry when coupled with UPLC can be a highly selective and sensitive method for quantitatively measuring analytes in complex matrices but is still subject to interferences and suppression. Additionally, by utilizing the Multiple Reaction Monitoring (MRM) feature of the triple quadrupole mass spectrometer, the signal-to-noise ratio can be significantly improved.

For this assay, the mass spectrometer was operated in positive ion mode with a voltage of 4000V. The source was heated to 300°C. Qaudrupole 1 (Q1) resolution was set to unit resolution with Quadrupole 3 (Q3) set to unit. Ion Source gas 1 & 2 and Curtain gas were all set to 40 psi. The CAD gas was set to 9. The MRM transitions monitored were for dA:

252.1→136.0 m/z with a Collision Energy (CE) of 28V and Collision Cell Potential (CXP) of 16V; for the IS: 267.1→146.0 m/z with a Collision Energy (CE) of 27V and Collision Cell Potential (CXP) of 7V; for m6A: 266.1→150.0 m/z with a Collision Energy (CE) of 22V and Collision Cell Potential (CXP) of 9V. The resulting chromatograms were integrated using Sciex OS: Autopeak software. Concentrations were calculated from a standard curve prepared in water.

*Inserts that were cloned into pMCs-Puro Retroviral Backbones*

pMCS -AID-mCherry-SV40-Puro Insert
atgaaggagaagagtgcttgtcctaaagatccagccaaacctccggccaaggcacaagttgtgggatggccaccggtgagatcata
ccggaagaacgtgatggtttcctgccaaaaatcaagcggtggcccggaggcggcggcgttcgtgaaggtatcaatggacggagca
ccgtacttgaggaaaatcgatttgaggatgtataaaatggtgagcaagggcgaggaggataacatggccatcatcaaggagttcatg
cgcttcaaggtgcacatggagggctccgtgaacggccacgagttcgagatcgagggcgagggcgagggccgcccctacgaggg
cacccagaccgccaagctgaaggtgaccaagggtggcccccctgcccttcgcctgggacatcctgtcccctcagttcatgtacggctcc
aaggcctacgtgaagcaccccgccgacatccccgactacttgaagctgtccttccccgagggcttcaagtgggagcgcgtgatgaa
cttcgaggacggcggcgtggtgaccgtgacccaggactcctccctgcaggacggcgagttcatctacaaggtgaagctgcgcggca
ccaacttcccctccgacggccccgtaatgcagaagaagaccatgggctgggaggcctcctccgagcggatgtaccccgaggacgg
cgccctgaagggcgagatcaagcagaggctgaagctgaaggacggcggccactacgacgctgaggtcaagaccacctacaag
gccaagaagcccgtgcagctgcccggcgcctacaacgtcaacatcaagttggacatcacctcccacaacgaggactacaccatcg
tggaacagtacgaacgcgccgagggccgccactccaccggcggcatggacgagctgtacaagtag

pMCS-UNG2-AID-mCherry-SV40-Puro Insert
atgatcggccagaagacgctctactcctttttctcccccagccccgccaggaagcgacacgcccccagccccgagccggccgtcca
ggggaccggcgtggctggggtgcctgaggaaagcggagatgcggcggccatcccagccaagaaggccccggctgggcaggag
gagcctgggacgccgccctcctcgccgctgagtgccgagcagttggaccggatccagaggaacaaggccgcggccctgctcaga
ctcgcggcccgcaacgtgcccgtgggctttggagagagctggaagaagcacctcagcggggagttcgggaaaccgtattttatcaa
gctaatgggatttgttgcagaagaaagaaagcattacactgtttatccacccccacaccaagtcttcacctggacccagatgtgtgaca
taaaagatgtgaaggttgtcatcctgggacaggatccatatcatggacctaatcaagctcacgggctctgctttagtgttcaaaggcctg
ttccgcctccgcccagtttggagaacatttataaagagttgtctacagacatagaggattttgttcatcctggccatggagatttatctgggt
gggccaagcaaggtgttctccttctcaacgctgtcctcacggttcgtgcccatcaagccaactctcataaggagcgaggctgggagca
gttcactgatgcagttgtgtcctggctaaatcagaactcgaatggccttgttttcttgctctggggctcttatgctcagaagaagggcagtgc
cattgataggaagcggcaccatgtactacagacggctcatccctcccctttgtcagtgtatagagggttctttggatgtagacacttttcaa
agaccaatgagctgctgcagaagtctggcaagaagcccattgactggaaggagctgaaggagaagagtgcttgtcctaaagatcc
agccaaacctccggccaaggcacaagttgtgggatggccaccggtgagatcataccggaagaacgtgatggtttcctgccaaaaat
caagcggtggcccggaggcggcggcgttcgtgaaggtatcaatggacggagcaccgtacttgaggaaaatcgatttgaggatgtat
aaaatggtgagcaagggcgaggaggataacatggccatcatcaaggagttcatgcgcttcaaggtgcacatggagggctccgtga
acggccacgagttcgagatcgagggcgagggcgagggccgcccctacgagggcacccagaccgccaagctgaaggtgaccaa
gggtggcccccctgcccttcgcctgggacatcctgtcccctcagttcatgtacggctccaaggcctacgtgaagcaccccgccgacatc
cccgactacttgaagctgtccttccccgagggcttcaagtgggagcgcgtgatgaacttcgaggacggcggcgtggtgaccgtgacc
caggactcctccctgcaggacggcgagttcatctacaaggtgaagctgcgcggcaccaacttcccctccgacggccccgtaatgca
gaagaagaccatgggctgggaggcctcctccgagcggatgtaccccgaggacggcgccctgaagggcgagatcaagcagagg
ctgaagctgaaggacggcggccactacgacgctgaggtcaagaccacctacaaggccaagaagcccgtgcagctgcccggcgc
ctacaacgtcaacatcaagttggacatcacctcccacaacgaggactacaccatcgtggaacagtacgaacgcgccgagggccg
ccactccaccggcggcatggacgagctgtacaagtag

*Inserts that were cloned into pLenti-CMV-Insert-SV40-Puro Backbones*

pLenti-CMV-Cry2-mCherry-SV40-Puro Insert
atgaagatggacaaaaagactatagtttggtttagaagagacctaaggattgaggataatcctgcattagcagcagctgctcacgaa
ggatctgtttttcctgtcttcatttggtgtcctgaagaagaaggacagttttatcctggaagagcttcaagatggtggatgaaacaatcactt
gctcacttatctcaatccttgaaggctcttggatctgacctcactttaatcaaaacccacaacacgatttcagcgatcttggattgtatccgc
gttaccggtgctacaaaagtcgtctttaaccacctctatgatcctgtttcgttagttcgggaccataccgtaaaggagaagctggtggaac
gtgggatctctgtgcaaagctacaatggagatctattgtatgaaccgtgggagatatactgcgaaaagggcaaacctttttacgagtttca
attcttactggaagaaatgcttagatatgtcgattgaatccgttatgcttcctcctccttggcggttgatgccaataactgcagcggctgaag
cgatttgggcgtgttcgattgaagaactagggctggagaatgaggccgagaaaccgagcaatgcgttgttaactagagcttggtctcc
aggatggagcaatgctgataagttactaaatgagttcatcgagaagcagttgatagattatgcaaagaacagcaagaaagttgttgg
gaattctacttcactactttctccgtatctccatttcggggaaataagcgtcagacacgttttccagtgtgcccggatgaaacaaattatatg
ggcaagagataagaacagtgaaggagaagaaagtgcagatctttttcttaggggaatcggtttaagagagtattctcggtatatatgttt
caacttcccgtttactcacgagcaatcgttgttgagtcatcttcggtttttcccttgggatgctgatgttgataagttcaaggcctggagacaa
ggcaggaccggttatccgttggtggatgccggaatgagagagctttgggctaccggatggatgcataacagaataagagtgattgttt
caagctttgctgtgaagtttcttctccttccatggaaatggggaatgaagtatttctgggatacacttttggatgctgatttggaatgtgacatc
cttggctggcagtatatctctgggagtatccccgatggccacgagcttgatcgcttggacaatcccgcgttacaaggcgccaaatatga
cccagaaggtgagtacataaggcaatggcttcccgagcttgcgagattgccaactgaatggatccatcatccatggacgctccttta
accgtactcaaagcttctggtgtggaactcggaacaaactatgcgaaacccattgtagacatcgacacagctcgtgagctactagcta
aagctatttcaagaacccgtggagcacagatcatgatcggagcagcagcccgggatccaccggtcgccaccatggtgagcaagg
gcgaggaggataacatggccatcatcaaggagttcatgcgcttcaaggtgcacatggagggctccgtgaacggccacgagttcgag
atcgagggcgagggcgagggccgcccctacgagggcacccagaccgccaagctgaaggtgaccaagggtggccccctgccctt
cgcctgggacatcctgtcccctcagttcatgtacggctccaaggcctacgtgaagcaccccgccgacatccccgactacttgaagctg
tccttccccgagggcttcaagtgggagcgcgtgatgaacttcgaggacggcggcgtggtgaccgtgacccaggactcctccctgcag
gacggcgagttcatctacaaggtgaagctgcgcggcaccaacttcccctccgacggccccgtaatgcagaagaagaccatgggct
gggaggcctcctccgagcggatgtaccccgaggacggcgccctgaagggcgagatcaagcagaggctgaagctgaaggacgg
cggccactacgacgctgaggtcaagaccacctacaaggccaagaagcccgtgcagctgcccggcgcctacaacgtcaacatcaa
gttggacatcacctcccacaacgaggactacaccatcgtggaacagtacgaacgcgccgagggccgccactccaccggcggcat
ggacgagctgtacaagtaa

pLenti-CMV-Cry2-mCherry-UNG2-SV40-Puro Insert
atgaagatggacaaaaagactatagtttggtttagaagagacctaaggattgaggataatcctgcattagcagcagctgctcacgaa
ggatctgtttttcctgtcttcatttggtgtcctgaagaagaaggacagttttatcctggaagagcttcaagatggtggatgaaacaatcactt
gctcacttatctcaatccttgaaggctcttggatctgacctcactttaatcaaaacccacaacacgatttcagcgatcttggattgtatccgc
gttaccggtgctacaaaagtcgtctttaaccacctctatgatcctgtttcgttagttcgggaccataccgtaaaggagaagctggtggaac
gtgggatctctgtgcaaagctacaatggagatctattgtatgaaccgtgggagatatactgcgaaaagggcaaacctttttacgagtttca
attcttactggaagaaatgcttagatatgtcgattgaatccgttatgcttcctcctccttggcggttgatgccaataactgcagcggctgaag
cgatttgggcgtgttcgattgaagaactagggctggagaatgaggccgagaaaccgagcaatgcgttgttaactagagcttggtctcc
aggatggagcaatgctgataagttactaaatgagttcatcgagaagcagttgatagattatgcaaagaacagcaagaaagttgttgg
gaattctacttcactactttctccgtatctccatttcggggaaataagcgtcagacacgttttccagtgtgcccggatgaaacaaattatatg
ggcaagagataagaacagtgaaggagaagaaagtgcagatctttttcttaggggaatcggtttaagagagtattctcggtatatatgttt
caacttcccgtttactcacgagcaatcgttgttgagtcatcttcggtttttcccttgggatgctgatgttgataagttcaaggcctggagacaa
ggcaggaccggttatccgttggtggatgccggaatgagagagctttgggctaccggatggatgcataacagaataagagtgattgttt
caagctttgctgtgaagtttcttctccttccatggaaatggggaatgaagtatttctgggatacacttttggatgctgatttggaatgtgacatc
cttggctggcagtatatctctgggagtatccccgatggccacgagcttgatcgcttggacaatcccgcgttacaaggcgccaaatatga
cccagaaggtgagtacataaggcaatggcttcccgagcttgcgagattgccaactgaatggatccatcatccatggacgctccttta
accgtactcaaagcttctggtgtggaactcggaacaaactatgcgaaacccattgtagacatcgacacagctcgtgagctactagcta
aagctatttcaagaacccgtggagcacagatcatgatcggagcagcagcccgggatccaccggtcgccaccatggtgagcaagg
gcgaggaggataacatggccatcatcaaggagttcatgcgcttcaaggtgcacatggagggctccgtgaacggccacgagttcgag
atcgagggcgagggcgagggccgcccctacgagggcacccagaccgccaagctgaaggtgaccaagggtggccccctgccctt
cgcctgggacatcctgtcccctcagttcatgtacggctccaaggcctacgtgaagcaccccgccgacatccccgactacttgaagctg
tccttccccgagggcttcaagtgggagcgcgtgatgaacttcgaggacggcggcgtggtgaccgtgacccaggactcctccctgcag

gacggcgagttcatctacaaggtgaagctgcgcggcaccaacttcccctccgacggccccgtaatgcagaagaagaccatgggct
gggaggcctcctccgagcggatgtaccccgaggacggcgccctgaagggcgagatcaagcagaggctgaagctgaaggacgg
cggccactacgacgctgaggtcaagaccacctacaaggccaagaagcccgtgcagctgcccggcgcctacaacgtcaacatcaa
gttggacatcacctcccacaacgaggactacaccatcgtggaacagtacgaacgcgccgagggccgccactccaccggcggcat
ggacgagctgtacaagatgatcggccagaagacgctctactcctttttctcccccagccccgccaggaagcgacacgcccccagcc
ccgagccggccgtccaggggaccggcgtggctggggtgcctgaggaaagcggagatgcggcggccatcccagccaagaaggc
cccggctgggcaggaggagcctgggacgccgccctcctcgccgctgagtgccgagcagttggaccggatccagaggaacaagg
ccgcggccctgctcagactcgcggcccgcaacgtgcccgtgggctttggagagagctggaagaagcacctcagcggggagttcgg
gaaaccgtattttatcaagctaatgggatttgttgcagaagaaagaaagcattacactgtttatccacccccacaccaagtcttcacctg
gacccagatgtgtgacataaaagatgtgaaggttgtcatcctgggacaggatccatatcatggacctaatcaagctcacgggctctgct
ttagtgttcaaaggcctgttccgcctccgcccagtttggagaacatttataaagagttgtctacagacatagaggattttgttcatcctggcc
atggagatttatctgggtgggccaagcaaggtgttctccttctcaacgctgtcctcacggttcgtgcccatcaagccaactctcataagga
gcgaggctgggagcagttcactgatgcagttgtgtcctggctaaatcagaactcgaatggccttgttttcttgctctggggctcttatgctca
gaagaagggcagtgccattgataggaagcggcaccatgtactacagacggctcatccctcccctttgtcagtgtatagagggttctttg
gatgtagacacttttcaaagaccaatgagctgctgcagaagtctggcaagaagcccattgactggaaggagctgtaa

pLenti-CMV-Cry2-mCherry-IDR-SV40-Puro Insert
atgaagatggacaaaaagactatagtttggtttagaagagacctaaggattgaggataatcctgcattagcagcagctgctcacgaa
ggatctgtttttcctgtcttcatttggtgtcctgaagaagaaggacagttttatcctggaagagcttcaagatggtggatgaaacaatcactt
gctcacttatctcaatccttgaaggctcttggatctgacctcactttaatcaaaacccacaacacgatttcagcgatcttggattgtatccgc
gttaccggtgctacaaaagtcgtctttaaccacctctatgatcctgtttcgttagttcgggaccataccgtaaaggagaagctggtggaac
gtgggatctctgtgcaaagctacaatggagatctattgtatgaaccgtgggagatatactgcgaaaagggcaaaccttttacgagtttca
attcttactggaagaaatgcttagatatgtcgattgaatccgttatgcttcctcctccttggcggttgatgccaataactgcagcggctgaag
cgatttgggcgtgttcgattgaagaactagggctggagaatgaggccgagaaaccgagcaatgcgttgttaactagagcttggtctcc
aggatggagcaatgctgataagttactaaatgagttcatcgagaagcagttgatagattatgcaaagaacagcaagaaagttgttgg
gaattctacttcactactttctccgtatctccatttcggggaaataagcgtcagacacgttttccagtgtgcccggatgaaacaaattatatg
ggcaagagataagaacagtgaaggagaagaaagtgcagatctttttcttaggggaatcggtttaagagagtattctcggtatatatgttt
caacttcccgtttactcacgagcaatcgttgttgagtcatcttcggtttttcccttgggatgctgatgttgataagttcaaggcctggagacaa
ggcaggaccggttatccgttggtggatgccggaatgagagagctttgggctaccggatggatgcataacagaataagagtgattgttt
caagctttgctgtgaagtttcttctccttccatggaaatggggaatgaagtatttctgggatacacttttggatgctgatttggaatgtgacatc
cttggctggcagtatatctctgggagtatccccgatggccacgagcttgatcgcttggacaatcccgcgttacaaggcgccaaatatga
cccagaaggtgagtacataaggcaatggcttcccgagcttgcgagattgccaactgaatggatccatcatccatgggacgctccttta
accgtactcaaagcttctggtgtggaactcggaacaaactatgcgaaacccattgtagacatcgacacagctcgtgagctactagcta
aagctatttcaagaacccgtggagcacagatcatgatcggagcagcagcccgggatccaccggtcgccaccatggtgagcaagg
gcgaggaggataacatggccatcatcaaggagttcatgcgcttcaaggtgcacatggagggctccgtgaacggccacgagttcgag
atcgagggcgagggcgagggccgcccctacgagggcacccagaccgccaagctgaaggtgaccaagggtggccccctgccctt
cgcctgggacatcctgtcccctcagttcatgtacggctccaaggcctacgtgaagcaccccgccgacatccccgactacttgaagctg
tccttccccgagggcttcaagtgggagcgcgtgatgaacttcgaggacggcggcgtggtgaccgtgacccaggactcctccctgcag
gacggcgagttcatctacaaggtgaagctgcgcggcaccaacttcccctccgacggccccgtaatgcagaagaagaccatgggct
gggaggcctcctccgagcggatgtaccccgaggacggcgccctgaagggcgagatcaagcagaggctgaagctgaaggacgg
cggccactacgacgctgaggtcaagaccacctacaaggccaagaagcccgtgcagctgcccggcgcctacaacgtcaacatcaa
gttggacatcacctcccacaacgaggactacaccatcgtggaacagtacgaacgcgccgagggccgccactccaccggcggcat
ggacgagctgtacaagatgatcggccagaagacgctctactcctttttctcccccagccccgccaggaagcgacacgcccccagcc
ccgagccggccgtccaggggaccggcgtggctggggtgcctgaggaaagcggagatgcggcggccatcccagccaagaaggc
cccggctgggcaggaggagcctgggacgccgccctcctcgccgctgagtgccgagcagttggaccggatccagaggaacaagg
ccgcggccctgctcagactcgcggcccgcaacgtgtaa

pLenti-CMV-Cry2-mCherry-ΔIDR -SV40-Puro Insert
atgaagatggacaaaaagactatagtttggtttagaagagacctaaggattgaggataatcctgcattagcagcagctgctcacgaa
ggatctgtttttcctgtcttcatttggtgtcctgaagaagaaggacagttttatcctggaagagcttcaagatggtggatgaaacaatcactt
gctcacttatctcaatccttgaaggctcttggatctgacctcactttaatcaaaacccacaacacgatttcagcgatcttggattgtatccgc

gttaccggtgctacaaaagtcgtctttaaccacctctatgatcctgtttcgttagttcgggaccataccgtaaaggagaagctggtggaac
gtgggatctctgtgcaaagctacaatggagatctattgtatgaaccgtgggagatatactgcgaaaagggcaaaccttttacgagtttca
attcttactggaagaaatgcttagatatgtcgattgaatccgttatgcttcctcctccttggcggttgatgccaataactgcagcggctgaag
cgatttgggcgtgttcgattgaagaactagggctggagaatgaggccgagaaaccgagcaatgcgttgttaactagagcttggtctcc
aggatggagcaatgctgataagttactaaatgagttcatcgagaagcagttgatagattatgcaaagaacagcaagaaagttgttgg
gaattctacttcactactttctccgtatctccatttcggggaaataagcgtcagacacgttttccagtgtgcccggatgaaacaaattatatg
ggcaagagataagaacagtgaaggagaagaaagtgcagatctttttcttaggggaatcggtttaagagagtattctcggtatatatgttt
caacttcccgtttactcacgagcaatcgttgttgagtcatcttcggtttttcccttgggatgctgatgttgataagttcaaggcctggagacaa
ggcaggaccggttatccgttggtggatgccggaatgagagagctttgggctaccggatggatgcataacagaataagagtgattgttt
caagctttgctgtgaagtttcttctccttccatggaaatggggaatgaagtatttctgggatacacttttggatgctgatttggaatgtgacatc
cttggctggcagtatatctctgggagtatccccgatggccacgagcttgatcgcttggacaatcccgcgttacaaggcgccaaatatga
cccagaaggtgagtacataaggcaatggcttcccgagcttgcgagattgccaactgaatggatccatcatccatgggacgctccttta
accgtactcaaagcttctggtgtggaactcggaacaaactatgcgaaacccattgtagacatcgacacagctcgtgagctactagcta
aagctatttcaagaacccgtggagcacagatcatgatcggagcagcagcccgggatccaccggtcgccaccatggtgagcaagg
gcgaggaggataacatggccatcatcaaggagttcatgcgcttcaaggtgcacatggagggctccgtgaacggccacgagttcgag
atcgagggcgagggcgagggccgcccctacgagggcacccagaccgccaagctgaaggtgaccaagggtggcccctgccctt
cgcctgggacatcctgtcccctcagttcatgtacggctccaaggcctacgtgaagcaccccgccgacatccccgactacttgaagctg
tccttccccgagggcttcaagtgggagcgcgtgatgaacttcgaggacggcggcgtggtgaccgtgacccaggactcctccctgcag
gacggcgagttcatctacaaggtgaagctgcgcggcaccaacttcccctccgacggccccgtaatgcagaagaagaccatgggct
gggaggcctcctccgagcggatgtaccccgaggacggcgccctgaagggcgagatcaagcagaggctgaagctgaaggacgg
cggccactacgacgctgaggtcaagaccacctacaaggccaagaagcccgtgcagctgcccggcgcctacaacgtcaacatcaa
gttggacatcacctcccacaacgaggactacaccatcgtggaacagtacgaacgcgccgagggccgccactccaccggcggcat
ggacgagctgtacaagcccgtgggctttggagagagctggaagaagcacctcagcggggagttcgggaaaccgtattttatcaagc
taatgggatttgttgcagaagaaagaaagcattacactgtttatccacccccacaccaagtcttcacctggacccagatgtgtgacata
aaagatgtgaaggttgtcatcctgggacaggatccatatcatggacctaatcaagctcacgggctctgctttagtgttcaaaggcctgttc
cgcctccgcccagtttggagaacatttataaagagttgtctacagacatagaggattttgttcatcctggccatggagatttatctgggtgg
gccaagcaaggtgttctccttctcaacgctgtcctcacggttcgtgcccatcaagccaactctcataaggagcgaggctgggagcagtt
cactgatgcagttgtgtcctggctaaatcagaactcgaatggccttgttttcttgctctggggctcttatgctcagaagaagggcagtgcca
ttgataggaagcggcaccatgtactacagacggctcatccctcccctttgtcagtgtatagagggttctttggatgtagacacttttcaaag
accaatgagctgctgcagaagtctggcaagaagcccattgactggaaggagctgtaa

pLenti-CMV-Cry2-mCherry-IDR-C-SV40-Puro Insert
atgaagatggacaaaaagactatagtttggtttagaagagacctaaggattgaggataatcctgcattagcagcagctgctcacgaa
ggatctgtttttcctgtcttcatttggtgtcctgaagaagaaggacagttttatcctggaagagcttcaagatggtggatgaaacaatcactt
gctcacttatctcaatccttgaaggctcttggatctgacctcactttaatcaaaacccacaacacgatttcagcgatcttggattgtatccgc
gttaccggtgctacaaaagtcgtctttaaccacctctatgatcctgtttcgttagttcgggaccataccgtaaaggagaagctggtggaac
gtgggatctctgtgcaaagctacaatggagatctattgtatgaaccgtgggagatatactgcgaaaagggcaaaccttttacgagtttca
attcttactggaagaaatgcttagatatgtcgattgaatccgttatgcttcctcctccttggcggttgatgccaataactgcagcggctgaag
cgatttgggcgtgttcgattgaagaactagggctggagaatgaggccgagaaaccgagcaatgcgttgttaactagagcttggtctcc
aggatggagcaatgctgataagttactaaatgagttcatcgagaagcagttgatagattatgcaaagaacagcaagaaagttgttgg
gaattctacttcactactttctccgtatctccatttcggggaaataagcgtcagacacgttttccagtgtgcccggatgaaacaaattatatg
ggcaagagataagaacagtgaaggagaagaaagtgcagatctttttcttaggggaatcggtttaagagagtattctcggtatatatgttt
caacttcccgtttactcacgagcaatcgttgttgagtcatcttcggtttttcccttgggatgctgatgttgataagttcaaggcctggagacaa
ggcaggaccggttatccgttggtggatgccggaatgagagagctttgggctaccggatggatgcataacagaataagagtgattgttt
caagctttgctgtgaagtttcttctccttccatggaaatggggaatgaagtatttctgggatacacttttggatgctgatttggaatgtgacatc
cttggctggcagtatatctctgggagtatccccgatggccacgagcttgatcgcttggacaatcccgcgttacaaggcgccaaatatga
cccagaaggtgagtacataaggcaatggcttcccgagcttgcgagattgccaactgaatggatccatcatccatgggacgctccttta
accgtactcaaagcttctggtgtggaactcggaacaaactatgcgaaacccattgtagacatcgacacagctcgtgagctactagcta
aagctatttcaagaacccgtggagcacagatcatgatcggagcagcagcccgggatccaccggtcgccaccatggtgagcaagg
gcgaggaggataacatggccatcatcaaggagttcatgcgcttcaaggtgcacatggagggctccgtgaacggccacgagttcgag
atcgagggcgagggcgagggccgcccctacgagggcacccagaccgccaagctgaaggtgaccaagggtggcccctgccctt

cgcctgggacatcctgtcccctcagttcatgtacggctccaaggcctacgtgaagcaccccgccgacatccccgactacttgaagctg
tccttccccgagggcttcaagtgggagcgcgtgatgaacttcgaggacggcggcgtggtgaccgtgacccaggactcctccctgcag
gacggcgagttcatctacaaggtgaagctgcgcggcaccaacttcccctccgacggccccgtaatgcagaagaagaccatgggct
gggaggcctcctccgagcggatgtaccccgaggacggcgccctgaagggcgagatcaagcagaggctgaagctgaaggacgg
cggccactacgacgctgaggtcaagaccacctacaaggccaagaagcccgtgcagctgcccggcgcctacaacgtcaacatcaa
gttggacatcacctcccacaacgaggactacaccatcgtggaacagtacgaacgcgccgagggccgccactccaccggcggcat
ggacgagctgtacaagcccgtgggctttggagagagctggaagaagcacctcagcggggagttcgggaaaccgtattttatcaagc
taatgggatttgttgcagaagaaagaaagcattacactgtttatccacccccacaccaagtcttcacctggacccagatgtgtgacata
aaagatgtgaaggttgtcatcctgggacaggatccatatcatggacctaatcaagctcacgggctctgctttagtgttcaaaggcctgttc
cgcctccgcccagtttggagaacatttataaagagttgtctacagacatagaggattttgttcatcctggccatggagatttatctgggtgg
gccaagcaaggtgttctccttctcaacgctgtcctcacggttcgtgcccatcaagccaactctcataaggagcgaggctgggagcagtt
cactgatgcagttgtgtcctggctaaatcagaactcgaatggccttgttttcttgctctggggctcttatgctcagaagaagggcagtgcca
ttgataggaagcggcaccatgtactacagacggctcatccctcccctttgtcagtgtatagagggttctttggatgtagacacttttcaaag
accaatgagctgctgcagaagtctggcaagaagcccattgactggaaggagctgatcggccagaagacgctctactcctttttctccc
ccagccccgccaggaagcgacacgcccccagccccgagccggccgtccaggggaccggcgtggctggggtgcctgaggaaag
cggagatgcggcggccatcccagccaagaaggccccggctgggcaggaggagcctgggacgccgccctcctcgccgctgagtg
ccgagcagttggaccggatccagaggaacaaggccgcggccctgctcagactcgcggcccgcaacgtgtaa

pLenti-CMV-Cry2-mCherry-ΔPIP-SV40-Puro Insert
atgaagatggacaaaaagactatagtttggtttagaagagacctaaggattgaggataatcctgcattagcagcagctgctcacgaa
ggatctgtttttcctgtcttcatttggtgtcctgaagaagaaggacagttttatcctggaagagcttcaagatggtggatgaaacaatcactt
gctcacttatctcaatccttgaaggctcttggatctgacctcactttaatcaaaacccacaacacgatttcagcgatcttggattgtatccgc
gttaccggtgctacaaaagtcgtctttaaccacctctatgatcctgtttcgttagttcgggaccataccgtaaaggagaagctggtggaac
gtgggatctctgtgcaaagctacaatggagatctattgtatgaaccgtgggagatatactgcgaaaagggcaaaccttttacgagtttca
attcttactggaagaaatgcttagatatgtcgattgaatccgttatgcttcctcctccttggcggttgatgccaataactgcagcggctgaag
cgatttgggcgtgttcgattgaagaactagggctggagaatgaggccgagaaaccgagcaatgcgttgttaactagagcttggtctcc
aggatggagcaatgctgataagttactaaatgagttcatcgagaagcagttgatagattatgcaaagaacagcaagaaagttgttgg
gaattctacttcactactttctccgtatctccatttcggggaaataagcgtcagacacgttttccagtgtgcccggatgaaacaaattatatg
ggcaagagataagaacagtgaaggagaagaaagtgcagatctttttcttaggggaatcggtttaagagagtattctcggtatatatgttt
caacttcccgtttactcacgagcaatcgttgttgagtcatcttcggtttttcccttgggatgctgatgttgataagttcaaggcctggagacaa
ggcaggaccggttatccgttggtggatgccggaatgagagagctttgggctaccggatggatgcataacagaataagagtgattgttt
caagctttgctgtgaagtttcttctccttccatggaaatggggaatgaagtatttctgggatacacttttggatgctgatttggaatgtgacatc
cttggctggcagtatatctctgggagtatccccgatggccacgagcttgatcgcttggacaatcccgcgttacaaggcgccaaatatga
cccagaaggtgagtacataaggcaatggcttcccgagcttgcgagattgccaactgaatggatccatcatccatgggacgctccttta
accgtactcaaagcttctggtgtggaactcggaacaaactatgcgaaacccattgtagacatcgacacagctcgtgagctactagcta
aagctatttcaagaacccgtggagcacagatcatgatcggagcagcagcccgggatccaccggtcgccaccatggtgagcaagg
gcgaggaggataacatggccatcatcaaggagttcatgcgcttcaaggtgcacatggagggctccgtgaacggccacgagttcgag
atcgagggcgagggcgagggccgcccctacgagggcacccagaccgccaagctgaaggtgaccaagggtggccccctgccctt
cgcctgggacatcctgtcccctcagttcatgtacggctccaaggcctacgtgaagcaccccgccgacatccccgactacttgaagctg
tccttccccgagggcttcaagtgggagcgcgtgatgaacttcgaggacggcggcgtggtgaccgtgacccaggactcctccctgcag
gacggcgagttcatctacaaggtgaagctgcgcggcaccaacttcccctccgacggccccgtaatgcagaagaagaccatgggct
gggaggcctcctccgagcggatgtaccccgaggacggcgccctgaagggcgagatcaagcagaggctgaagctgaaggacgg
cggccactacgacgctgaggtcaagaccacctacaaggccaagaagcccgtgcagctgcccggcgcctacaacgtcaacatcaa
gttggacatcacctcccacaacgaggactacaccatcgtggaacagtacgaacgcgccgagggccgccactccaccggcggcat
ggacgagctgtacaagAtgatcggccagaagacgctctactccgcagcatcccccagccccgccaggaagcgacacgcccccca
gccccgagccggccgtccaggggaccggcgtggctggggtgcctgaggaaagcggagatgcggcggccatcccagccaagaa
ggccccggctgggcaggaggagcctgggacgccgccctcctcgccgctgagtgccgagcagttggaccggatccagaggaaca
aggccgcggccctgctcagactcgcggcccgcaacgtgcccgtgggctttggagagagctggaagaagcacctcagcggggagtt
cgggaaaccgtattttatcaagctaatgggatttgttgcagaagaaagaaagcattacactgtttatccacccccacaccaagtcttcac
ctggacccagatgtgtgacataaaagatgtgaaggttgtcatcctgggacaggatccatatcatggacctaatcaagctcacgggctct
gctttagtgttcaaaggcctgttccgcctccgcccagtttggagaacatttataaagagttgtctacagacatagaggattttgttcatcctg

gccatggagatttatctgggtgggccaagcaaggtgttctccttctcaacgctgtcctcacggttcgtgcccatcaagccaactctcataa
ggagcgaggctgggagcagttcactgatgcagttgtgtcctggctaaatcagaactcgaatggccttgttttcttgctctggggctcttatg
ctcagaagaagggcagtgccattgataggaagcggcaccatgtactacagacggctcatccctcccctttgtcagtgtatagagggtt
ctttggatgtagacacttttcaaagaccaatgagctgctgcagaagtctggcaagaagcccattgactggaaggagctgtaa

pLenti-CMV-Cry2-mCherry-DRPA-SV40-Puro Insert
atgaagatggacaaaaagactatagtttggtttagaagagacctaaggattgaggataatcctgcattagcagcagctgctcacgaa
ggatctgtttttcctgtcttcatttggtgtcctgaagaagaaggacagttttatcctggaagagcttcaagatggtggatgaaacaatcactt
gctcacttatctcaatccttgaaggctcttggatctgacctcactttaatcaaaacccacaacacgatttcagcgatcttggattgtatccgc
gttaccggtgctacaaaagtcgtctttaaccacctctatgatcctgtttcgttagttcgggaccataccgtaaaggagaagctggtggaac
gtgggatctctgtgcaaagctacaatggagatctattgtatgaaccgtgggagatatactgcgaaaagggcaaaccttttacgagtttca
attcttactggaagaaatgcttagatatgtcgattgaatccgttatgcttcctcctccttggcggttgatgccaataactgcagcggctgaag
cgatttgggcgtgttcgattgaagaactagggctggagaatgaggccgagaaaccgagcaatgcgttgttaactagagcttggtctcc
aggatggagcaatgctgataagttactaaatgagttcatcgagaagcagttgatagattatgcaaagaacagcaagaaagttgttgg
gaattctacttcactactttctccgtatctccatttcggggaaataagcgtcagacacgttttccagtgtgcccggatgaaacaaattatatg
ggcaagagataagaacagtgaaggagaagaaagtgcagatctttttcttaggggaatcggtttaagagagtattctcggtatatatgttt
caacttcccgtttactcacgagcaatcgttgttgagtcatcttcggtttttcccttgggatgctgatgttgataagttcaaggcctggagacaa
ggcaggaccggttatccgttggtggatgccggaatgagagagctttgggctaccggatggatgcataacagaataagagtgattgttt
caagctttgctgtgaagtttcttctccttccatggaaatggggaatgaagtatttctgggatacacttttggatgctgatttggaatgtgacatc
cttggctggcagtatatctctgggagtatccccgatggccacgagcttgatcgcttggacaatcccgcgttacaaggcgccaaatatga
cccagaaggtgagtacataaggcaatggcttcccgagcttgcgagattgccaactgaatggatccatcatccatgggacgctccttta
accgtactcaaagcttctggtgtggaactcggaacaaactatgcgaaacccattgtagacatcgacacagctcgtgagctactagcta
aagctatttcaagaacccgtggagcacagatcatgatcggagcagcagcccgggatccaccggtcgccaccatggtgagcaagg
gcgaggaggataacatggccatcatcaaggagttcatgcgcttcaaggtgcacatggagggctccgtgaacggccacgagttcgag
atcgagggcgagggcgagggccgcccctacgagggcacccagaccgccaagctgaaggtgaccaagggtggccccctgccctt
cgcctgggacatcctgtcccctcagttcatgtacggctccaaggcctacgtgaagcaccccgccgacatccccgactacttgaagctg
tccttccccgagggcttcaagtgggagcgcgtgatgaacttcgaggacggcggcgtggtgaccgtgacccaggactcctccctgcag
gacggcgagttcatctacaaggtgaagctgcgcggcaccaacttcccctccgacggccccgtaatgcagaagaagaccatgggct
gggaggcctcctccgagcggatgtaccccgaggacggcgccctgaagggcgagatcaagcagaggctgaagctgaaggacgg
cggccactacgacgctgaggtcaagaccacctacaaggccaagaagcccgtgcagctgcccggcgcctacaacgtcaacatcaa
gttggacatcacctcccacaacgaggactacaccatcgtggaacagtacgaacgcgccgagggccgccactccaccggcggcat
ggacgagctgtacaagAtgatcggccagaagacgctctactcctttttctcccccagccccgccaggaagcgacacgcccccagcc
ccgagccggccgtccaggggaccggcgtggctggggtgcctgaggaaagcggagatgcggcggccatcccagccaagaaggc
cccggctgggcaggaggagcctgggacgccgccctcctcgccgctgagtgccgagcagttggaccggatccagaggaacaagg
ccgcggccctgctcagactcgcggcctgtaacgtgcccgtgggctttggagagagctggaagaagcacctcagcggggagttcggg
aaaccgtattttatcaagctaatgggatttgttgcagaagaaagaaagcattacactgtttatccacccccacaccaagtcttcacctgg
acccagatgtgtgacataaaagatgtgaaggttgtcatcctgggacaggatccatatcatggacctaatcaagctcacgggctctgcttt
agtgttcaaaggcctgttccgcctccgcccagtttggagaacatttataaagagttgtctacagacatagaggattttgttcatcctggcca
tggagatttatctgggtgggccaagcaaggtgttctccttctcaacgctgtcctcacggttcgtgcccatcaagccaactctcataaggag
cgaggctgggagcagttcactgatgcagttgtgtcctggctaaatcagaactcgaatggccttgttttcttgctctggggctcttatgctcag
aagaagggcagtgccattgataggaagcggcaccatgtactacagacggctcatccctcccctttgtcagtgtatagagggttctttgg
atgtagacacttttcaaagaccaatgagctgctgcagaagtctggcaagaagcccattgactggaaggagctgtaa

### KEY RESOURCES TABLE

| REAGENT or RESOURCE | SOURCE | IDENTIFIER |
|---|---|---|
| Antibodies | | |
| Anti-N6-methyladenosine (anti-6mA) | Synaptic System | Catalog # 202 003, RRID: AB_2279214 |

| | | |
|---|---|---|
| Anti-phospho-Histone H2A.X (Ser139). Clone JBW301 | Millipore | 05-636; RRID: AB_309864 |
| Anti-mCherry (chicken) | Abcam | Catalog # Ab205402, RRID: AB_2722769 |
| Anti-mCherry (rabbit) | Abcam | Catalog # Ab213511; RRID: AB_2814891 |
| Alexa Fluor 488-conjugated Anti-mouse IgG (H+L) | Invitrogen | Catalog # A11029 |
| Alexan Fluor 488-conjugated anti-Chicken IgY (H+L) | Invitrogen | Catalog # A32931 |
| Anti-UNG | AbClonal | Catalog # A1261 (WB: 1:1000); RRID: AB_2759453 |
| Anti-METTL3 | AbClonal | Catalog # 8370 (WB: 1:1000); RRID: AB_2770344 |
| Anti-Tubulin, Clone DM1A (mouse) | Millipore | Catalog # MABT205; RRID: AB_11204167 |
| Anti-WTAP (rabbit) | Bethyl | Catalog # A301-435A; RRID: AB_961137 |
| IRDye 680RD Goat anti-rabbit | Licor | Catalog # 926-68071 |
| IRDye 800CW Donkey anti-mouse | Licor | Catalog # 926-32212 |
| IgG | Abcam | Catalog # Ab172730, RRID_2687931 |
| UNG | Abclonal | Catalog # A1261; RRID: AB_2759453 |
| Chemicals, Peptides, and Recombinant Proteins | | |
| Phosphate Buffered Saline (PBS) | Corning | Catalog # 21-040-CV |
| Heat Inactivated Fetal Bovine Serum (FBS) | Gibco | Catalog # 16140-071 |
| RPMI-1640 | Corning | Catalog # 10-040-CM |
| McCoy’s 5A | Gibco | Catalog # 16600-108 |
| DMEM | Gibco | Catalog # 11995073 |
| DMEM (No Phenol Red) | Gibco | Catalog # A1443001 |
| Penicillin Streptomycin Solution, 100x | Corning | Catalog # 30-002-CI |
| Penicillin Streptomycin Solution | Gibco | Catalog # 15120-122; used for U2OS 2-6-3 |
| GlutaMax | Gibco | Catalog # 35050061 |
| 0.25% Trypsin | Corning | Catalog # 25-053-CI |
| Recovery Cell Culture Freezing Medium | Gibco | Catalog # 12648010 |
| HAT Supplement | Gibco | Catalog # 21060-017 |
| Floxuridine | Sigma Aldrich | Catalog # F0503 |
| Raltitrexed | Sigma Aldrich | Catalog # R9156 |
| Gemcitabine | Sigma Aldrich | Catalog # G6423 |
| Hydroxyurea | Usp | Catalog # 1332000 |
| Mitomycin C | StemCell Technologies | Catalog # 73273 |
| METTL3 inhibitor | MedChem Express | Catalog # HY-134836/CS-0159584 |
| Puromycin | ThermoScientific | Catalog # J67236.XF |
| Hygromycin B | ThermoFisher | Catalog # 10687010 |
| 6-Thioguanine | Tocris | Catalog # 4061 |
| pCMV-Gag-Pol | CellBioLabs | Catalog #: RV-111 |
| pCMV-VSV-G | CellBioLabs | Catalog #: RV-110 |

| | | |
|---|---|---|
| Lentiviral Packaging Construct Mix | Sigma | Catalog # SHP001 |
| OPTI-MEM | Gibco | Catalog # 31985-062 |
| Polybrene Transfection Reagent | Millipore | Catalog # TR-10030G |
| Puromycin | InvivoGen | Catalog # Ant-pr |
| Lipofectamine™ 3000 Transfection Reagent | Invitrogen | Catalog # L300075 |
| Protein G Dynabeads | Invitrogen | Catalog # 10004D |
| TCEP Bond Breaker | ThermoFisher | Catalog # 77720 |
| Halt Protease & Phosphatase Inhibitor | ThermoFisher | Catalog # 78436 |
| Benzonase | Sigma Aldrich | Catalog # 70664-10KUN |
| Dithiothreitol (DTT) | Sigma Aldrich | Catalog # D0632-10G |
| Iodoacetamide (IAA) | Sigma Aldrich | Catalog # I1149-5G |
| Lysyl Endopeptidase (LysC) | Fujifilm Wako Chemicals USA | Catalog # 125-05061 |
| Formic acid | Fisher Chemical | Catalog # A117-50 |
| Sep-Pak C-18 | Waters | Catalog # WAT036925 |
| Trypsin | Promega | Catalog # V5111 |
| Easy-Spray 50cm column packed with 2 μm C-18 Resin | ThermoFisher | Catalog # ES903 |
| DNA Degradase Plus | Zymo Research | Catalog # 214843 |
| 10X DNA Degrader Reaction Buffer | Zymo Research | Catalog # E2016-2 |
| 2'-deoxyadenosine (dA) | Sigma | Catalog # D7400 |
| N6-methyl-2-deoxyadenosine (6mA) | ThermoFisher | Catalog #AAJ64961MD |
| stable heavy labeled 2'-deoxyadenosine | Cambridge Isotope Laboratories, Inc | CNLM-3896-CA-25; internal standard for analyte mass spectrometry |
| NuPAGE LDS Sample Buffer (4X) | Invitrogen | Catalog # NP0007 |
| NuPAGE Sample Reducing Agent (10X) | Invitrogen | Catalog # NP0009 |
| MOPS SDS Running Buffer (20X) | Invitrogen | Catalog # NP0001 |
| Invitrogen™ iBlot™ 2 Transfer Stacks, PVDF, mini | Invitrogen | Catalog # IB24002 |
| Chameleon Duo Prestained Protein ladder | LiCor | Catalog # 928-60000 |
| Tuberculin Needle | BD | |
| Intercept Blocking Buffer | LiCor | Catalog # 927-70001 |
| Phosphate Buffered Saline-Tween (20X) | Boston Bioproducts Inc | Catalog # IBB-920 |
| FBS | Gibco | Catalog # 16000-044 |
| 1 M HEPES | Corning | Catalog # 25-060-CI |
| 0.5 M EDTA | Invitrogen | Catalog # 46-000-CM |
| NaCl | Sigma | Catalog # S3014-1K |
| Triton X-100 | ThermoScientific | Catalog # A16046.AE |
| Sucrose | Thermo | Catalog # 036508.30 |
| $MgCl_2$ | Fluka | Catalog # 63020-1L |
| 37% Formaldehyde | ThermoScientific | Catalog # BP531-25 |
| DAPI | ThermoScientific | Catalog # 62248 (use at 1:10,000) |
| Duplex Buffer | IDT | Catalog # 1072570 |
| Electroporation Enhancer | IDT | Catalog # 1075916 |
| Alt-R CRISPR Cas9 tracrRNA | IDT | Catalog # 1073190 |
| Alt-R S.p. Cas9 Nuclease V3 | IDT | Catalog # 1081059 |
| Duplex Buffer | IDT | Catalog # 11-01-03-01 |
| Amaxa SE Cell Line Kit | Lonza | Catalog # V4SC-1096 |
| ---Solution Box | In Kit | Catalog # PBC1-02250 |
| ---SE solution | In Kit | Catalog # S-09637 |

| | | |
|---|---|---|
| ---Supplement Solution | In Kit | Catalog # S-09699 |
| TaqMan Gene Expression Master Mix | ThermoFisher | Catalog # 4369016 |
| Taqman Assay – GAPDH | ThermoFisher | Catalog # 4331182, Assay ID Hs99999905_m1 |
| Taqman Assay - UNG | ThermoFisher | Catalog # 4331182, Assay ID Hs01037093_m1, |
| DMEM for SILAC | ThermoFisher | Catalog # 88364 |
| $^{13}C_6$ L-Arginine-HCl | ThermoFisher | Catalog # 88210 |
| $^{13}C_6$ L-Lysine-2HCl | ThermoFisher | Catalog # 88209 |
| biotin-phenol | LGC GENOMICS LLC | Catalog # 41994-02-9 |
| doxycycline | Sigma Aldrich | Catalog # D9891-1G |
| $H_2O_2$ | Sigma Aldrich | Catalog # H1009 |
| sodium ascorbate | Sigma Aldrich | Catalog # A7631 |
| Trolox | Sigma Aldrich | Catalog # 238813 |
| sodium azide | Sigma Aldrich | Catalog # S2002 |
| Phosphate Buffered Saline (PBS) | Gibco | Catalog # 10010049 |
| cOmplete protease inhibitor cocktail | Sigma | Catalog # 11873580001 |
| Methanol | Sigma | Catalog # 1793307 |
| Crystal Violet | Aqua Solutions | Catalog # C8126 |
| Recombinant UNG-Catalytic Domain | This Paper | This Paper |
| Critical Commercial Assays | | |
| QIAquick PCR Cleanup Kit | Qiagen | Catalog # 28506 |
| PureLink Quick PCR Purification Kit | Invitrogen | Catalog # K310001 |
| Gentra Puregene kit | Qiagen | Catalog # 158845 |
| MTS Assay kit | Abcam | Catalog # ab197010 |
| Quick-DNA/RNA Miniprep Plus Kit | Zymo Research | Catalog # D7003 |
| RNeasy Plus University Kit | Qiagen | Catalog # 730404 |
| High Capacity RT Kit | Applied Biosystems | Catalog # 4374966 |
| Experimental Models: Cell Lines | | |
| Human: DLD-1 | ATCC | Catalog # CCL-221 |
| Human: DLD-1 UNG KO Clone E7 | This Paper | This Paper |
| Human: DLD-1 UNG KO Clone E7 + pMCS_UNG2_AID_mCherry | This Paper | This Paper |
| Human: DLD-1 UNG KO Clone E7 + pMCS_AID_mCherry | This Paper | This Paper |
| Human: DLD-1 UNG KO Clone E7 + pCMV_Cry2_mCherry_EV | This Paper | This Paper |
| Human: DLD-1 UNG KO Clone E7 + pCMV_Cry2_mCherry_UNG2 | This Paper | This Paper |
| Human: DLD-1 UNG KO Clone E7 + pCMV_Cry2_mCherry_UNG2_IDR | This Paper | This Paper |
| Human: DLD-1 UNG KO Clone E7 + pCMV_Cry2_mCherry_UNG2_ΔIDR | This Paper | This Paper |
| Human: DLD-1 UNG KO Clone E7 + pCMV_Cry2_mCherry_UNG2_IDR-C | This Paper | This Paper |
| Human: DLD-1 UNG KO Clone E7 + pCMV_Cry2_mCherry_UNG2_ ΔPIP | This Paper | This Paper |
| Human: DLD-1 UNG KO Clone E7 + pCMV_Cry2_mCherry_UNG2_ ΔRPA | This Paper | This Paper |
| Human: HT-29 | ATCC | Catalog # HTB-38 |
| Human: HT-29 pLenti7-EF1a-Cas9 | This Paper | This Paper |
| Human: SW620 | ATCC | Catalog # CCL-227 |
| Human: U2OS 2-6-3 | Spector Lab | PMID: 15006351 |

| | | |
|---|---|---|
| Human: U2OS 2-6-3 + GFP-LacI-APEX2-UNG2$^{IDR}$ | This Paper | This Paper |
| Oligonucleotides | | |
| gRNA_UNG_4,<br>CTTGATGGGCACGAACCGTG | IDT | N/A |
| gRNA_HPRT,<br>AATTATGGGGATTACTAGGA | IDT | N/A; targets intronic region |
| gRNA_METTL3-ex10-1,<br>CAGTTGGGTTGCACATTGTG | IDT | N/A |
| gRNA_UNG-2597,<br>TCCCCTTTGTCAGTGTATAG | IDT | N/A |
| gRNA_WTAP_Hs.Cas9.WTAP.1.AB | IDT | Catalog # 313817305 |
| gRNA_METTL3_ Hs.Cas9.METTL3.1.AA | IDT | Catalog # 313817302 |
| gRNA_NTC<br>GTAGCGAACGTGTCCGGCGT | IDT | N/A |
| dsDNA-U:A, 5' -<br>/5Biosg//iSp9/AAATTGUTATCCGCT<br>Complement:<br>5'-AGCGGATAACAATTT | IDT | N/A |
| dsDNA-U:m6dA, 5' -<br>/5Biosg//iSp9/AAATTGUTATCCGCT<br>Complement:<br>5'-<br>AGCGGATA/iN6Me-dA/CAATTT | IDT | N/A |
| dsDNA-U:A, m6dA:T, 5' -<br>/5Biosg//iSp9/AAATTGUT/iN6Me-dA/TCCGCT<br>Complement:<br>5'-AGCGGATAACAATTT | IDT | N/A |
| ssDNA-U:<br>/5Biosg//iSp9/AAATTGUTATCCGCT | IDT | N/A |
| ssDNA-U_m6dA:<br>/5Biosg//iSp9/AAATTGUT/iN6Me-dA/TCCGCT | IDT | N/A |
| Recombinant DNA | | |
| pLenti-EF1a-Cas9 | Pfizer | #5342 |
| Custom sgRNA library | DeskGen | This Paper |
| pMCS-Puro Retroviral Vector | CellBio Labs | Catalog # RTV-041 |
| pMCS_AID_mCherry_Puro Retroviral Vector | Azenta Life Sciences | N/A - C096 - See supplemental information |
| pMCS_UNG2_AID_mCherry Retroviral Vector | Azenta Life Sciences | N/A - C096 - See supplemental information |
| pLenti-GIII-CMV | Applied Biological Materials | 16422061 |
| pLenti-CMV-Cry2-mCherry-SV40-Puro (EV) Lentiviral Vector | Applied Biological Materials | N/A - C096 - See supplemental information |
| pLenti-CMV-Cry2-mCherry-UNG2-SV40-Puro Lentiviral Vector | Applied Biological Materials | N/A - C096 - See supplemental information |
| pLenti-CMV-Cry2-mCherry-UNG2-IDR-SV40-Puro Lentiviral Vector | Applied Biological Materials | N/A - C096 - See supplemental information |

| | | |
|---|---|---|
| pLenti-CMV-Cry2-mCherry-UNG2-ΔIDR-SV40-Puro Lentiviral Vector | Applied Biological Materials | N/A - C096 - See supplemental information |
| pLenti-CMV-Cry2-mCherry-UNG2-IDR-C-SV40-Puro Lentiviral Vector | Applied Biological Materials | N/A - C096 - See supplemental information |
| pLenti-CMV-Cry2-mCherry-UNG2-ΔPIP-SV40-Puro Lentiviral Vector | Applied Biological Materials | N/A - C096 - See supplemental information |
| pLenti-CMV-Cry2-mCherry- ΔRPA-SV40-Puro Lentiviral Vector | Applied Biological Materials | N/A - C096 - See supplemental information |
| Software and Algorithms | | |
| Model-based Analysis of Genome-wide CRISPR-Cas9 Knockout (MAGecK) | | MLE |
| ImageJ | | 1.47v |
| PRISM | Graph Pad Software | Version 9 |
| Sciex OS: Autopeak | Sciex | 2.2.0 |
| CellProfilier | | |
| MaxQuant | | 1.6.17.0 |
| Other | | |
| CX7 CellNightSight | ThermoFisher | Immunofluorescence |
| UltraView Spinning Disk | PerkinElmer | Cry2 Imaging |
| Incucyte | Sartorius | Cell Viability - Growth |
| Odyssey CX7 | Li-cor | Immunoblotting |
| Illumina Next-Seq | Illumina | Whole Genome Screen |
| Illumina Mi-Seq | Illumina | Whole Genome Screen |
| 4D Nucleofector | Lonza | KO line generation |
| Envision 2104 Plate Reader | Perkin Elmer | Cell Viability - MTS |
| nanoACQUITY UPLC® System | Waters | Coimmunoprecipitation LC-MS/MS |
| Orbitrap Fusion™ Lumos™ Tribrid™ Mass Spectrometer | ThermoFishser | Coimmunoprecipitation LC-MS/MS |
| ACQUITY UPLC® M Class System | Waters | UPLC-MS/MS |
| Triple Quad™ 7500 System | Sciex | UPLC-MS/MS |
| Column: nanoEase m/z peptide BEH c18, 300A, 1.7um 300um x 100mm | Waters | UPLC-MS/MS, PN186009264 |

**Figure S1. Generation of mCherry-tagged UNG KO DLD-1 cells.**

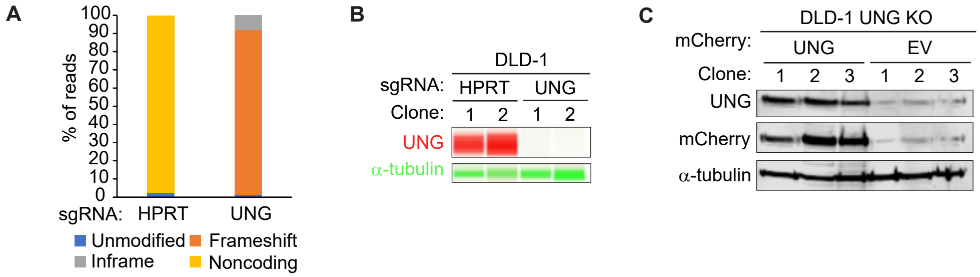

**Figure S2. UNG responds to uracil-based DNA damage in a manner partly dependent on its intrinsically disordered region.**

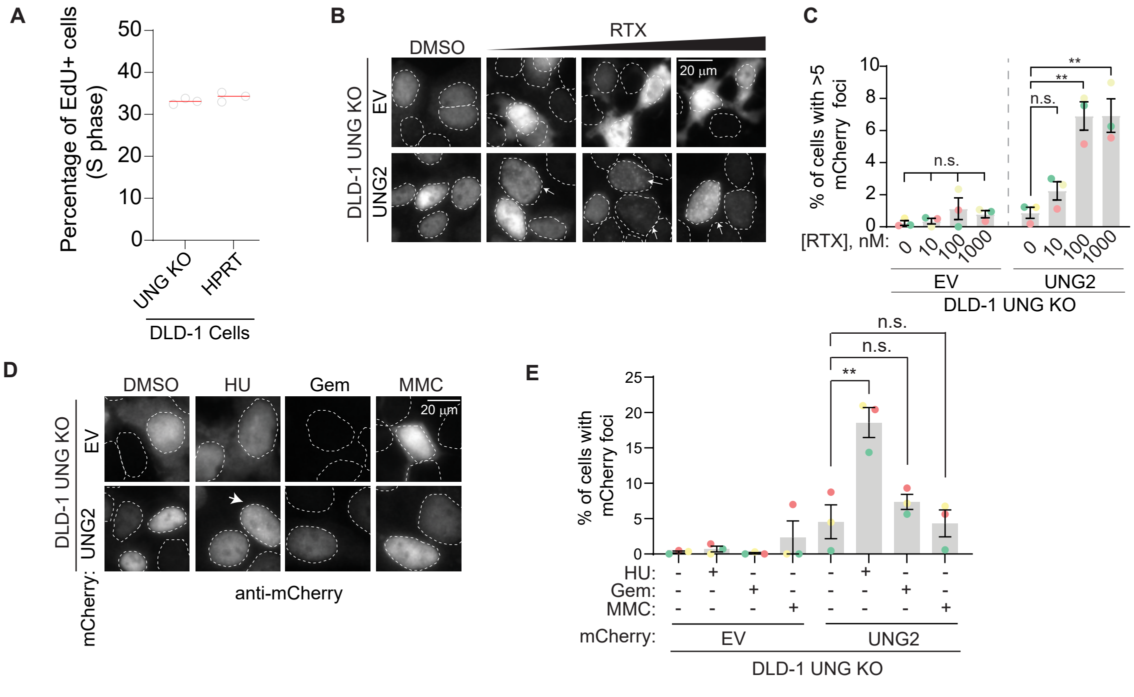


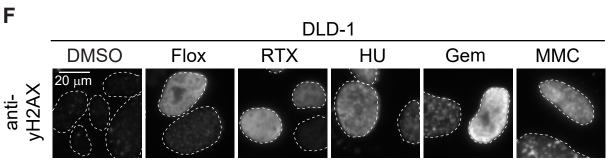


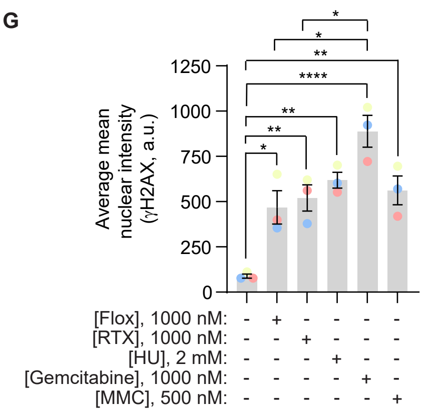


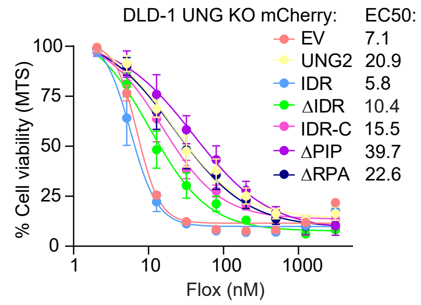


I

**KEGG Pathways**

| Description | Count in Network | FDR |
|---|---|---|
| Mismatch Repair | 15 of 23 | 2.09E-7 |
| Nucleotide Excision Reapir | 29 of 46 | 1.15E-13 |
| Spliceosome | 80 of 132 | 1.30E-37 |
| DNA Replication | 22 of 36 | 3.92E-10 |
| Base Excision Repair | 13 of 33 | 0.00015 |

**Figure S3. N6-methyladenosine foci in response to genomic uracil-inducing agents are not RNA:DNA hybrids.**

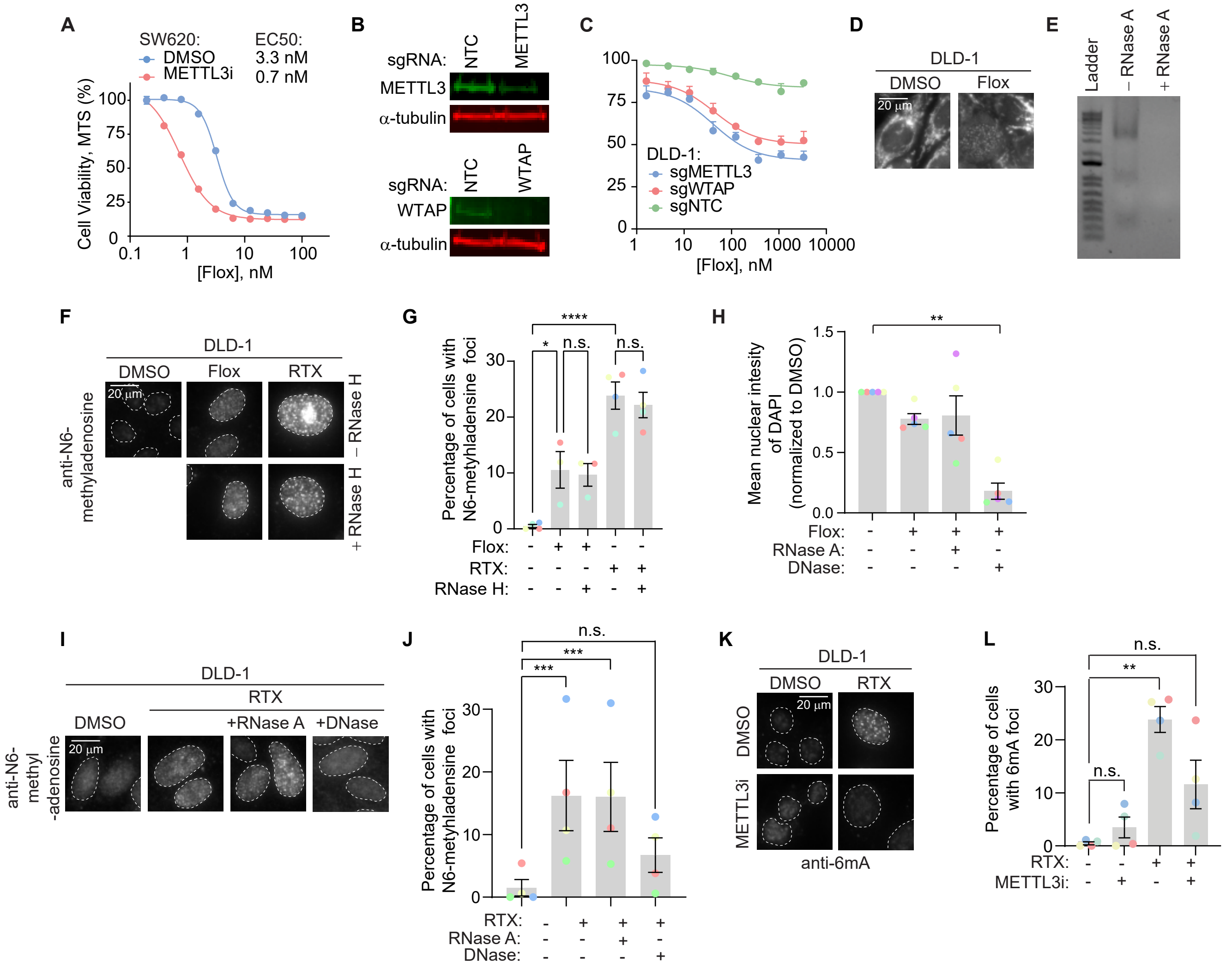

**Figure S4. The presence of 6mA does not alter UNG binding kinetics.**

**A**

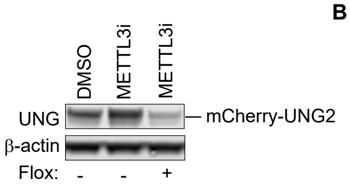


**B**

| Template | $K_D$ (pM) |
|---|---|
| dsDNA-U:A | 297 +/- 147 |
| dsDNA-U:6mA | 385 +/- 238 |
| dsDNA-U:A, 6mA:T | 361 +/- 207 |

**C**

| Template | $K_D$ (pM) |
|---|---|
| ssDNA-U | 103 +/- 22 |
| ssDNA-U, 6mA | 139 +/- 17 |

**Figure S5. 6mA foci, but not UNG foci, correlate with DNA damage levels.**

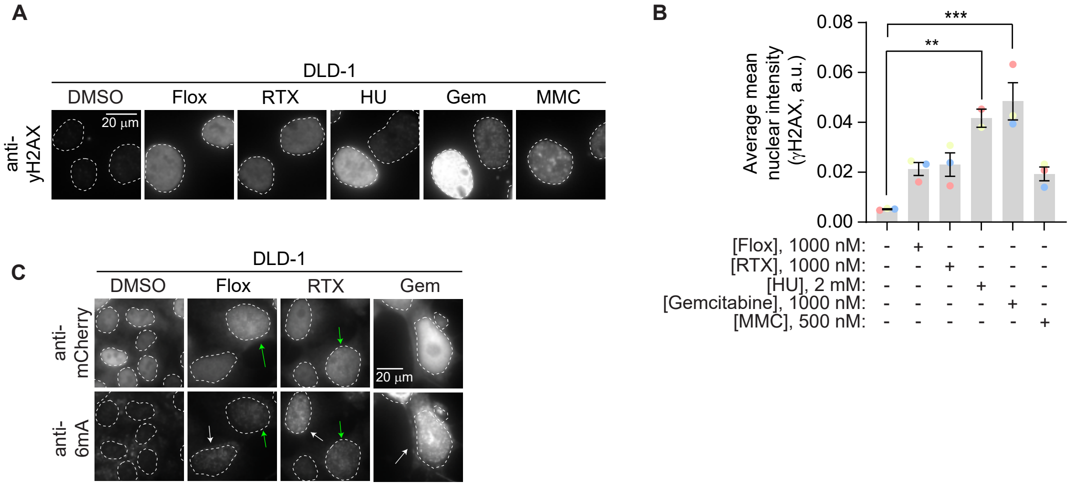
